# IRES-independent, eIF2A/eIF2D-dependent translation of the enterovirus genome

**DOI:** 10.1101/2022.06.01.494276

**Authors:** Hyejeong Kim, David Aponte-Diaz, Mohamad S. Sotoudegan, Jamie J. Arnold, Craig E. Cameron

## Abstract

RNA recombination in positive-strand RNA viruses is a molecular-genetic process, which permits the greatest evolution of the genome and may be essential to stabilizing the genome from the deleterious consequences of accumulated mutations. Enteroviruses represent a useful system to elucidate the details of this process. On the biochemical level, it is known that RNA recombination is catalyzed by the viral RNA-dependent RNA polymerase using a template-switching mechanism. For this mechanism to function in cells, the recombining genomes must be located in the same subcellular compartment. How a viral genome is trafficked to the site of genome replication and recombination, which is membrane associated and isolated from the cytoplasm, is not known. We hypothesized that genome translation was essential for co-localization of genomes for recombination. We show that complete inactivation of internal ribosome entry site (IRES)-mediated translation of a donor enteroviral genome enhanced recombination instead of impairing it. Recombination did not occur by a non-replicative mechanism. Rather, sufficient translation of the non-structural region of the genome occurred to support subsequent steps required for recombination. The non-canonical translation initiation factors, eIF2A and eIF2D, were required for IRES-independent translation. Our results support an eIF2A/eIF2D-dependent mechanism under conditions in which the eIF2-dependent mechanism is inactive. Detection of an IRES-independent mechanism for translation of the enterovirus genome provides an explanation for a variety of debated observations, including non-replicative recombination and persistence of enteroviral RNA lacking an IRES. The existence of an eIF2A/eIF2D-dependent mechanism in enteroviruses predicts the existence of similar mechanisms in other viruses.

## Introduction

RNA recombination is an important driver of evolution in positive-strand RNA viruses [1–3]. Myriad examples exist of the creation of pathogenic viral strains by RNA recombination [4–7]. Indeed, it has been suggested that RNA recombination contributed to the evolution of severe acute respiratory syndrome coronavirus 2 (SARS-CoV-2) and is contributing to creation of the ever-changing repertoire of variants circulating globally [8]. A second epidemiological challenge produced by RNA recombination is the creation of virulent, vaccine-derived polioviruses by recombination of live, attenuated vaccine strains with circulating, wild species- C enteroviruses [9]. In spite of the prevalence and importance of RNA recombination in virus evolution, an understanding of the biochemistry and cell biology of this process remains woefully incomplete.

Studies of RNA recombination in the species-C enterovirus, poliovirus, have established the molecular and conceptual framework guiding studies of RNA recombination in other positive-strand RNA viruses [10, 11]. Mixing of genetically encoded phenotypes between poliovirus variants occurs rapidly upon co-infection [12]. RNA recombination occurs by a template-switching mechanism in which the viral RNA-dependent RNA polymerase (RdRp) initiates RNA synthesis first on one genome (referred to as the donor template) then switches to a second genome (referred to as the acceptor template) during elongation [10, 13]. Transfection of two genomes, each of which individually incapable of producing infectious virus, uses RNA recombination to reconstitute an infectious genome [13]. Systems such as these have shown a direct correlation between genetic errors introduced into the genome and the frequency of recombination, implicating RdRp infidelity as a trigger of recombination [14–16].

The RdRp is sufficient for template switching in vitro [17]. Mutations mapping to RdRp-coding sequence can impair RNA recombination [18, 19]. These recombination-defective variants replicate well in cell culture under normal conditions [18, 19] but are unable to deal with high mutational loads [18] or replicate well in animals [18, 19]. Biophysical experiments revealed the existence of a stalled RdRp with its 3’-end of nascent RNA in a single-stranded form [20]. This state was inextricably linked to RNA recombination using a recombination-defective enzyme, thus identifying a putative recombination intermediate [19]. Formation of a single-stranded 3’-end while replicating the donor genome permits this end to hybridize to the acceptor genome [20]. Nucleotide misincorporation and incorporation of certain nucleotide analogs induces formation of the recombination intermediate, interfering with virus multiplication [19].

Enterovirus genome replication occurs in association with virus-induced membranes, which are also thought to be sequestered from the cytoplasm and mechanisms therein capable of sensing viral RNA [21]. This circumstance requires genomes undergoing recombination to co-localize to the same site of genome replication. How enteroviral genomes are targeted to sites of genome replication is not known. The initial goal of this study was to discover mechanisms governing genome trafficking.

We hypothesized that polyprotein determinants present on the translating ribosome have the capacity to target the translating polysome to the site of genome replication (**Fig. 1A**). Once present in the “genome-replication organelle,” translation would ultimately terminate because factors required for translation initiation would be present in the cytoplasm (**Fig. 1B**). These genomes would undergo replication and co-localization of distinct genomes would facilitate template switching and phenotypic mixing (**Fig. 1B**). If this hypothesis is correct, then interfering with translation of one of the genomes (**Fig. 1C**) would preclude trafficking to the genome-replication organelle and recombination (**Fig. 1D**).

**Fig 1.**
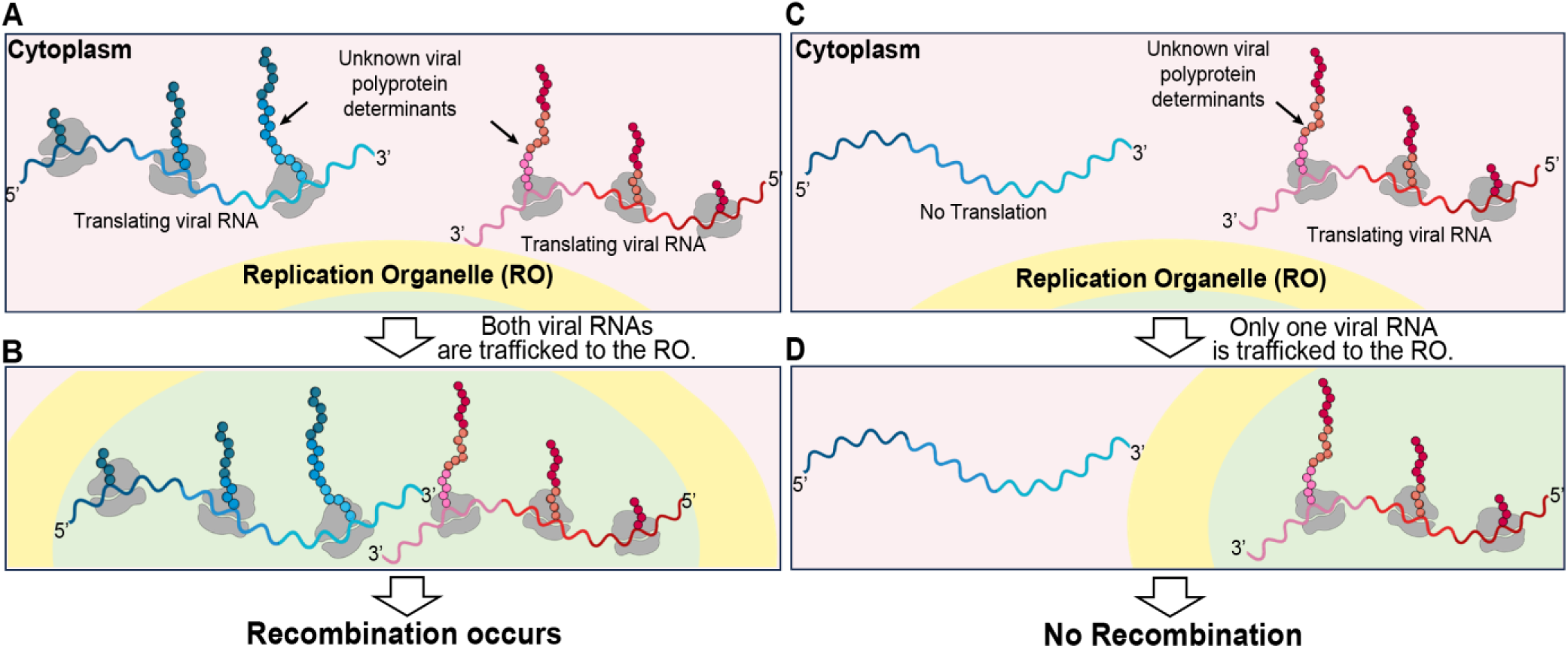
Hypothesis: Polyprotein determinants presented during translation direct the enteroviral genome to the site of replication. For recombination to occur between two genetically distinct PV genomes, these genomes must be in the same replication organelle. **(A, B)** Viral RNAs, shown in blue and red, are targeted to the replication organelle while being translated and facilitate RNA recombination between the two co-localized genomes. Specific viral polyprotein determinants, indicated by a black arrow, mediate this trafficking while still a part of the polyprotein and in association with the ribosome. **(C, D)** In a case where one genome with impaired polyprotein synthesis, shown in red, is coinfected in a cell with another genome, shown in blue, the genome with impaired polyprotein synthesis cannot be trafficked to the replication organelle so recombination between the two genomes cannot occur. This figure was created by Efraín E. Rivera-Serrano using BioRender.

The only known mechanism for translation of the enteroviral genome is the internal ribosome entry site (IRES) present in the 5’-untraslated region (UTR) of the genome [22–26]. Therefore, genetic inactivation of the IRES should impair recombination. This did not occur. Recombination was stimulated substantially. Some have suggested the existence of a replication-independent mechanism for recombination [27]. However, the corresponding study never investigated the possibility of an IRES-independent mechanism for translation. Herein we provide evidence for the existence of a translation-initiation mechanism that requires eIF2A and eIF2D, factors capable of initiating from non-AUG codons [28]. Our data are consistent with initiation occurring at the 3’-end of the 2A protease-coding sequence, downstream of the structural gene (P1) located at the 5’-end of the IRES-dependent open-reading frame (ORF). While population-level studies only revealed a role for eIF2A and eIF2D when in vitro transcribed RNA was transfected into cells to initiate infection and/or recombination, studies at the single-cell level suggested a role for these factors during normal, virus-initiated infection.

It is becoming increasingly clear that mechanisms exist for cells to survive under conditions of stress that lead to phosphorylation of eIF2α and shutdown of normal, cap-dependent translation [29]. One of these mechanisms is the use of non-AUG codons for translation initiation using eIF2A and/or eIF2D [30]. Among the earliest responses to viral infection is shutdown of cap-dependent translation by eIF2α kinases [31]. Our study suggests the very provocative possibility that enteroviruses may also exploit the eIF2A/eIF2D- dependent mechanism to survive under conditions in which the cell is actively attempting to thwart viral infection by inhibiting normal translation. Previous attempts to reveal a role for eIF2A and/or eIF2D in multiplication of other viruses may have been masked at the population level [32, 33]. Studies at the single-cell level may be necessary to uncover the importance of eIF2A and/or eIF2D in the viral lifecycle.

## Results

### Evidence for IRES-independent translation of the poliovirus genome

We have used the cell-based assay for poliovirus (PV) recombination developed by the Evans laboratory [13]. This assay employs two viral RNA genomes prepared in vitro by T7 transcription of the appropriate cDNAs [13]. The donor genome is a subgenomic replicon lacking capsid-coding sequence (donor in **Fig. 2A**). The acceptor genome is a complete PV genome; however, this genome is replication incompetent because of inactivating mutations within a cis-acting replication element (acceptor in **Fig. 2A**). Co-transfection of the donor and acceptor genomes into mammalian cells leads to the formation of recombinants capable of producing infectious virus (recombinant in **Fig. 2A**). Sufficient virus is produced by this system to permit virus titer to be used as a surrogate for recombination efficiency [13].

**Fig 2.**
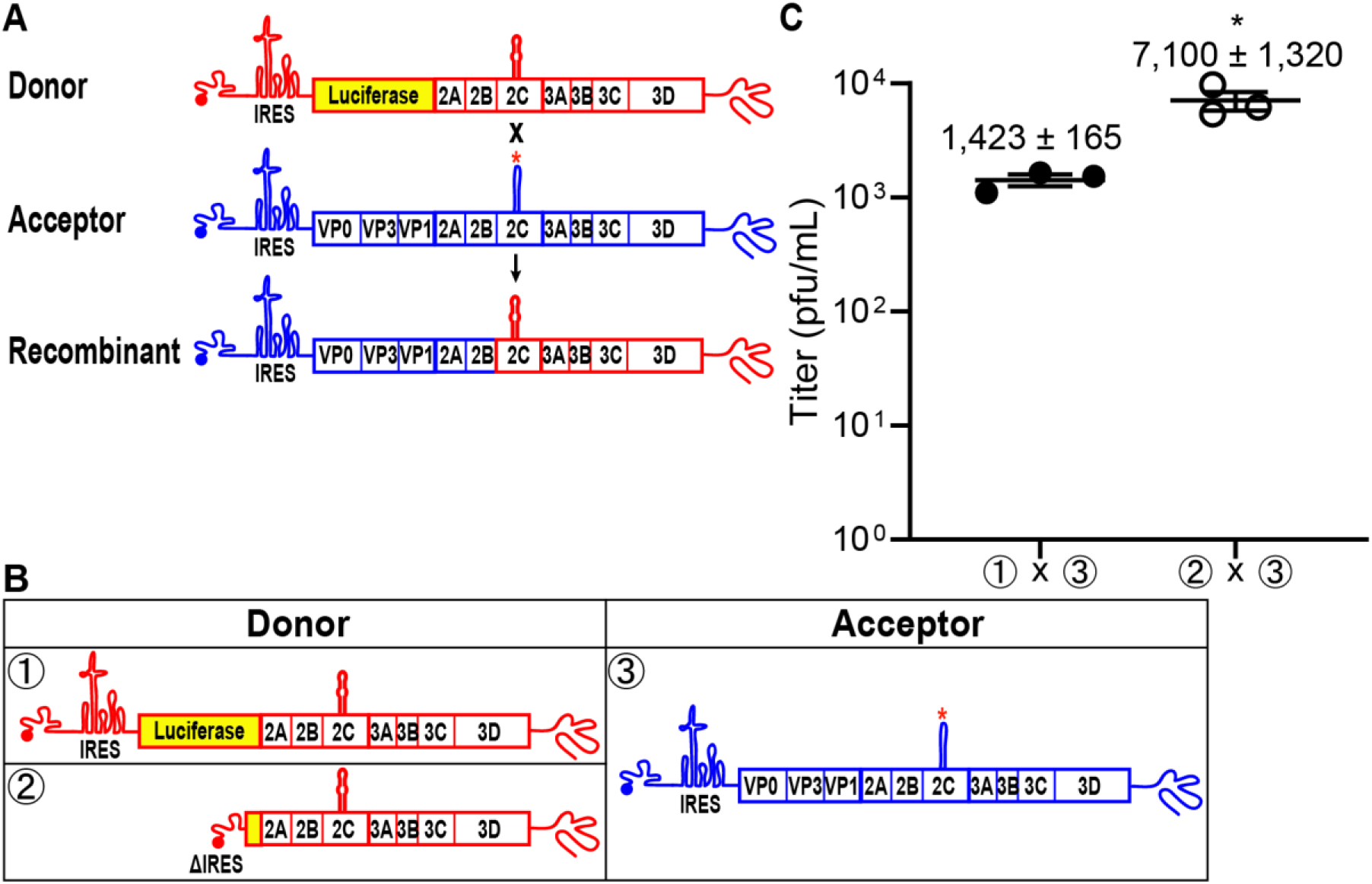
RNA virus recombinants are recovered after deletion of the PV IRES in a donor subgenomic replicon RNA. **(A)** Schematic of the cell-based assay for PV recombination [13, 14, 69]. Two RNAs are used in the assay, a replication-competent PV subgenomic RNA (Donor, red) in which capsid-coding sequence is replaced with luciferase coding-sequence and a replication-incompetent full-length PV genomic RNA (Acceptor, blue) with a defective cis-acting replication element (indicated by red *). Co-transfection of these RNAs in a L929 mouse fibroblast cell-line produces infectious virus if recombination occurs resulting in a replication-competent full-length PV genomic RNA. Infectious virus produced by recombination can be quantified by plaque assay using HeLa cells. **(B)** Comparison of infectious virus produced between donor RNAs with either an intact IRES (①) or when the entire IRES and majority of luciferase coding sequence (nt 41-2393, ΔIRES) was deleted (②) [13]. **(C)** Recombination between the donor with the deleted IRES (ΔIRES) and acceptor RNA (② x ③) produces five-fold more recombinant virus compared to the corresponding control (① x ③). Results show titer of recombinant virus (pfu/mL±SEM; *n = 3*). Statistical analyses were performed using unpaired, two-tailed t-test (*significance level *p* = < 0.05).

We have developed a standardized approach to depict and discuss the recombination experiments performed herein. Donor genomes will be drawn in red and numbered ①, ②, and so on (**Fig. 2B**). Acceptor genomes will be drawn in blue and numbering will pick up from where the donor numbering ended (e.g. ③ in **Fig. 2B**). Transfected donor-acceptor pairs will be labeled using the indicated genome numbering (e.g. ① х ③ in **Fig. 2C**).

If translation of both donor and acceptor genomes is required for localization within the same genome-replication organelle, then inactivation of translation on one of the genomes should impair recombination. To inactivate translation, we deleted the IRES in the donor template (② in **Fig. 2B**) and paired it with the standard acceptor genome (③ in **Fig. 2B**). In contrast to expectations, deletion of IRES enhanced recombination by 5-fold over the control (compare ② х ③ to ① х ③ in **Fig. 2C**).

Dogma in picornavirology has been that for a genome to be replicated, and therefore to serve as a template for recombination, translation of that genome must occur [34, 35]. However, it was possible that the polyprotein produced by the acceptor genome might complement the deficit of the donor genome. If this is the case, then eliminating production of the RdRp (3D) by the acceptor genome should inhibit recombination. To test this possibility, we constructed two acceptor genomes. We introduced two stop codons immediately following the 3B-coding sequence, which should eliminate expression of the 3C- and 3D-coding sequence (③ in **Fig. 3A**). Because it was unclear if the two stop codons would terminate translation with 100% efficiency, we constructed an acceptor genome that not only contained the two stop codons after 3B but also genetically inactivated the RdRp by mutating the sequence to alter its signature GDD motif to GAA (④ in **Fig. 3A**). Pairing the ΔIRES donor genome with either of the acceptor genomes still yielded recombinants (① х ③ and ① х ④ in **Fig. 3B**).

**Fig 3.**
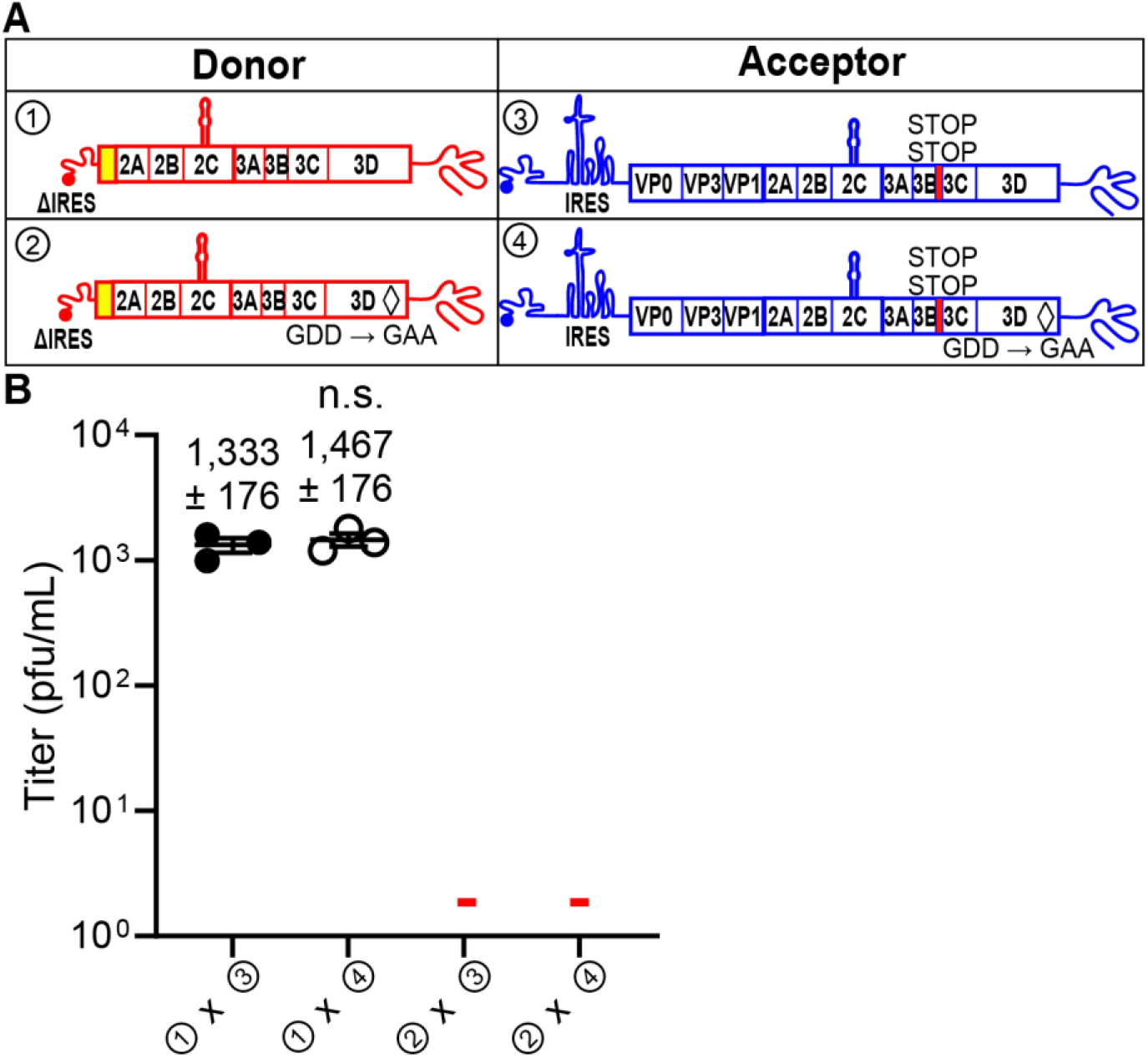
RNA virus recombinants are consistent with the donor RNA as the source of the PV RdRp, 3Dpol, and not the acceptor RNA. **(A)** A mutation (GDD to GAA) producing an inactive PV RdRp, shown by a black diamond (◊), was introduced into either the donor or acceptor RNAs (② and ④). Insertion of two STOP codons (UAGUAA) after the 3B-coding sequence (3B STOP), indicated by a red rectangle, was introduced into the acceptor RNAs (③ and ④). **(B)** The indicated donor and acceptor RNAs were co-transfected into L929 cells. Yields of recombinant virus following transfection are shown (pfu/mL±SEM; *n = 3*). **-** indicates that plaques were not detected (Limit of detection: 2 pfu/mL). Statistical analyses were performed using unpaired, two-tailed t-test (n.s. = non-significant). Viral recombinants were recovered only when the donor had an intact 3D gene encoding an active RdRp (① x ③, ① x ④); viral recombinants were not recovered when the donor RNA encoded an inactive RdRp (② x ③, ② x ④). There was no impact on viral recombinants produced when the acceptor RNA had stop codons upstream of the 3D gene encoding an active or inactive RdRp (① x ③, ① x ④).

At this stage, two possibilities remained to explain the observation of recombination using the ΔIRES donor genome. The first possibility was recombination by a non-replicative, RdRp-independent mechanism, and such a mechanism has been proposed to exist for enteroviruses [27]. However, the second possibility is the existence of an IRES-independent mechanism for translation of the PV genome. If this latter possibility were correct, then it would explain the observations that led to the existence of non-replicative recombination. By definition, a non-replicative mechanism should not require the RdRp. So, we constructed a ΔIRES donor genome that also harbored a genetically inactive RdRp (② in **Fig. 3A**). An inactive RdRp would be expected to only interfere with replicative recombination. Pairing the ΔIRES, inactive-RdRp donor genome with either of the acceptor genomes designed to preclude expression of the RdRp abrogated recombination (② х ③ and ② х ④ in **Fig. 3B**).

It now appeared that the donor genome was being translated. However, deleting the IRES creates a 5’-end essentially lacking a UTR. In this case, translation initiation might be an irrelevant mechanism. To probe this possibility, we inhibited IRES-dependent translation by deleting ten nucleotides from stem-loop II-3 (ΔSLII-3) in the donor genomes with or without an active RdRp; these constructs contained the entire luciferase-coding sequence (**Fig. 4A**) [36]. We paired each of the ΔSLII-3 donor genomes with each of the acceptor genomes designed to preclude expression of the RdRp. The outcomes with ΔSLII-3 donor genomes (**Fig. 4B**) were identical to those for the comparable ΔIRES donor genomes (**Fig. 3B**), consistent with the idea that an IRES-independent mechanism of translation exists for PV.

**Fig 4.**
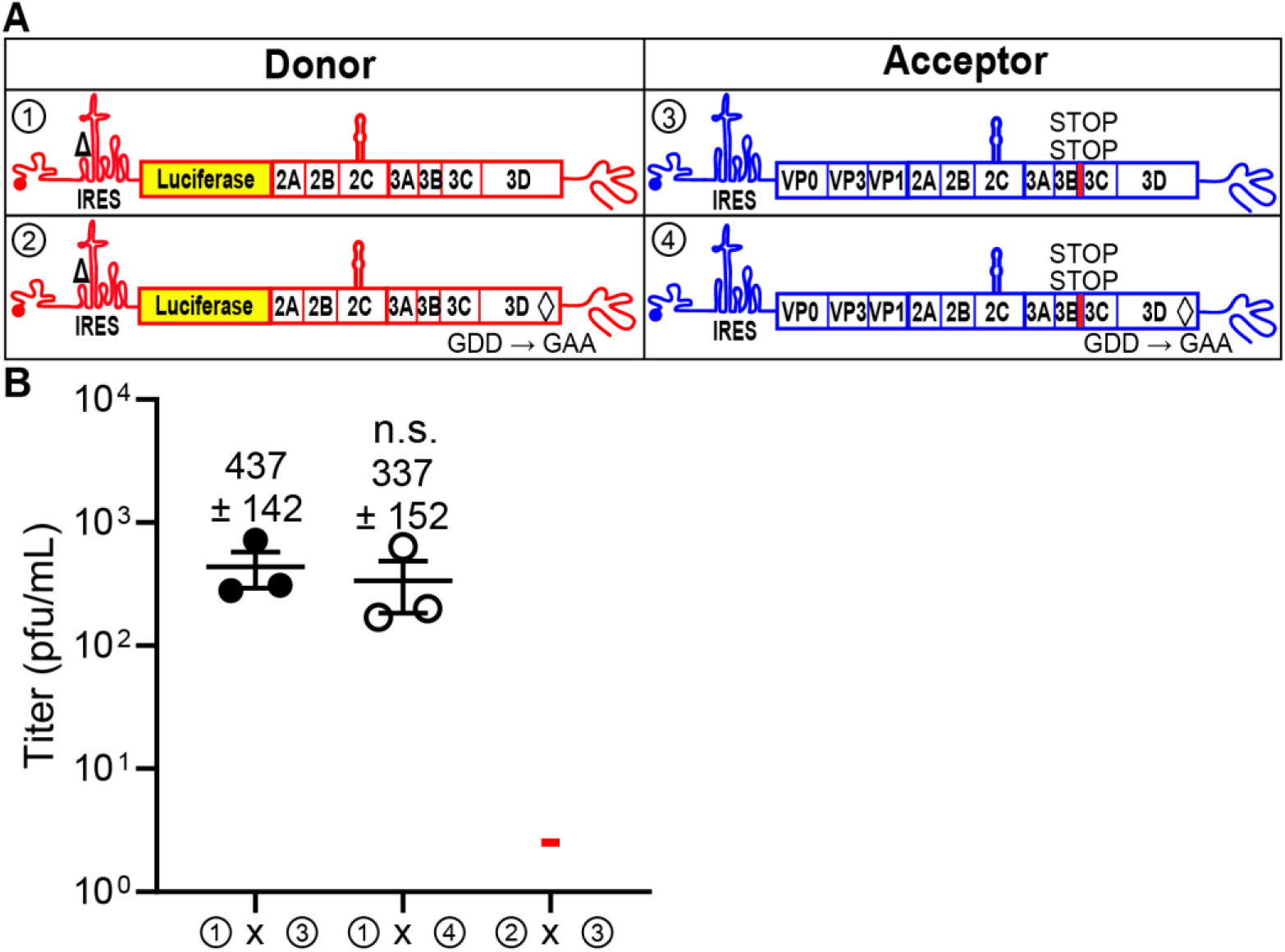
A 10-nt deletion in the IRES that is known to prevent translation recapitulates phenotypes observed with a deleted IRES. **(A)** The donor RNA was engineered to contain a ten-nucleotide deletion (nt 185-189, nt 198-202) known to disrupt the IRES (referred to as ΔSLII-3) [36] with an active or an inactive RdRp. **(B)** Comparison of infectious virus produced between the indicated donor and acceptor RNAs with the specific modifications. Results show titer of recombinant virus (pfu/mL±SEM; *n = 3*). **-** indicates that plaques were not detected (Limit of detection: 2 pfu/mL). Statistical analyses were performed using unpaired, two-tailed t-test (n.s. = non-significant). Viral recombinants were recovered using a donor RNA containing the ΔSLII-3 and an active RdRp with the indicated acceptor RNAs (① x ③, ① x ④); viral recombinants were not recovered when the donor RNA contained ΔSLII-3 and encoded an inactive RdRp (② x ③).

### Evidence for IRES-independent translation of the enterovirus A71 genome

It was important to evaluate this phenomenon of IRES-independent translation using another enterovirus, because conservation across the genus would be expected for a function essential to virus viability or fitness. We had already adapted EV-A71 for evaluation of recombination using the cell-based system [15]. We constructed an EV-A71 ΔIRES donor genome such that luciferase-coding sequence remained intact (**Fig. 5A**). Luciferase activity detected from the subgenomic RNA with a functional IRES in the presence of 3 mM guanidine hydrochloride (GuHCl), an inhibitor of enterovirus genome replication, showed accumulation of luciferase activity in cells (**Fig. 5A**). However, deletion of the IRES abolished accumulation of luciferase activity (**Fig. 5A**). Therefore, IRES-independent translation either produces luciferase at a level below the limit of detection or initiates downstream of luciferase-coding sequence.

**Fig 5.**
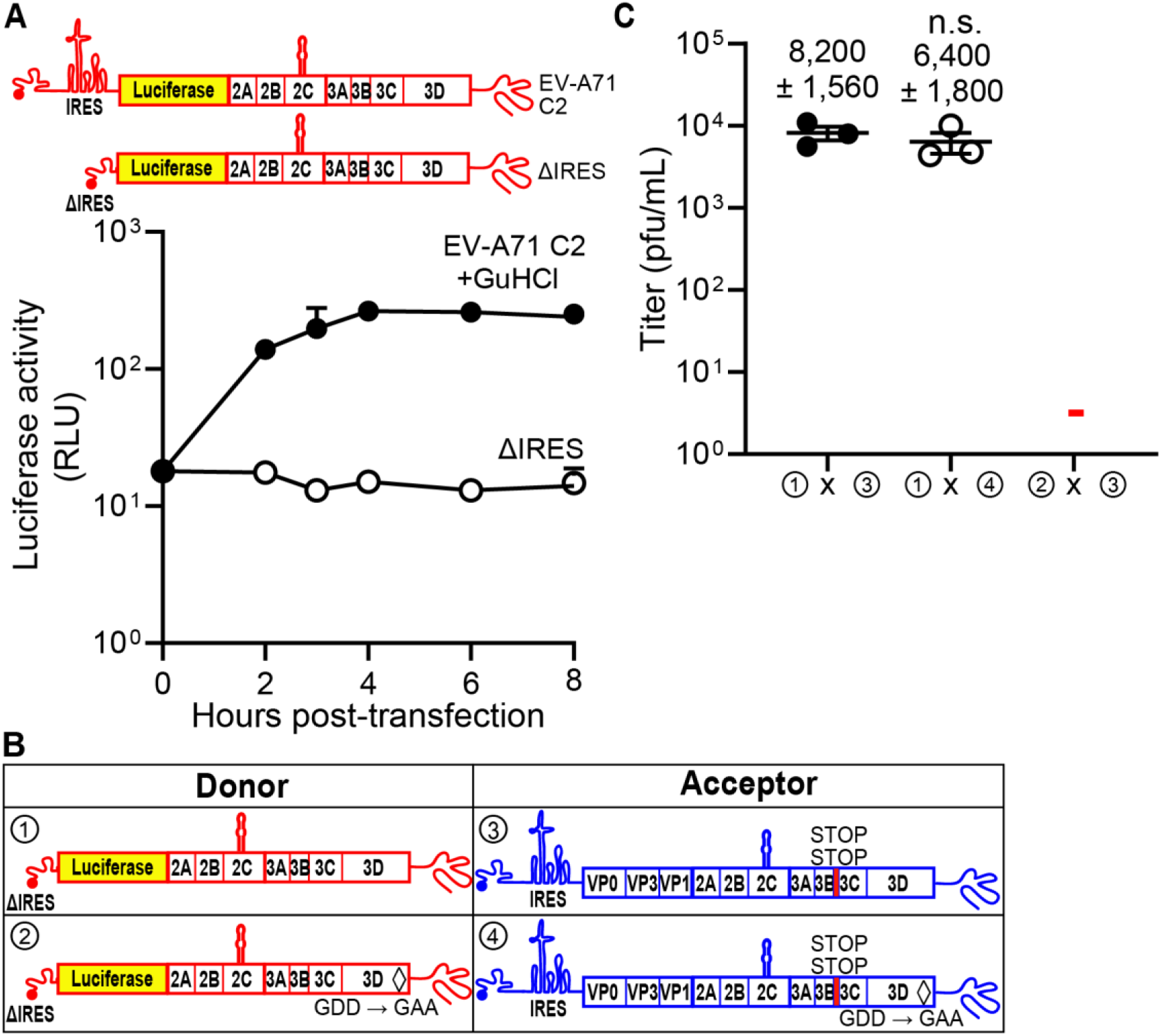
Evidence for IRES-independent translation of the enterovirus A71 genome. **(A)** Subgenomic replicon luciferase activity using an EV-A71 C2 subgenomic replicon with the entire IRES (nt 38-767) deleted (ΔIRES) [15]. As a control, the WT subgenomic replicon RNA was transfected in the presence of guanidine hydrochloride (GuHCl), a replication inhibitor. Luciferase activity is reported in relative light units (RLU) as a function of time post-transfection. **(B, C)** Comparison of infectious virus generated between the indicated donor and acceptor RNAs with the specific modifications. The indicated set of donor and acceptor RNAs was co-transfected into RD cells. Results show titer of recombinant virus (pfu/mL±SEM; *n = 3*). **-** indicates that plaques were not detected (Limit of detection: 2 pfu/mL). Statistical analyses were performed using unpaired, two-tailed t-test (n.s. = non-significant). Viral recombinants were recovered only when the donor had an intact 3D gene encoding an active RdRp (① x ③, ① x ④); viral recombinants were not recovered when the donor RNA encoded an inactive RdRp (② x ③).

We paired the EV-A71 ΔIRES donor genomes encoding an active or inactive RdRp with acceptor genomes incapable of expressing RdRp-coding sequence (**Fig. 5B**). As observed for PV (**Fig. 3B**), an active RdRp was required in the donor genome to observe recombination (**Fig. 5C**). These data are consistent with recombination occurring by a replicative, RdRp-dependent mechanism, with the RdRp being produced by an IRES-independent mechanism.

### IRES-independent translation may initiate within the non-structural protein-coding sequence

Studies above with EV-A71 donor genomes showed absolutely no detectable luciferase activity (**Fig. 5A**). We performed similar experiments for the PV genomes containing ΔIRES and deleted for luciferase-coding sequence (**Fig. 6A**) or ΔSLII-3 containing the complete luciferase-coding sequence (**Fig. 6B**). No detectable luciferase activity was observed for either construct with an inactive IRES (**Figs. 6A, B**). Indeed, the activity in the absence of a functional IRES was lower than the that detected for the corresponding WT donor genomes in the presence of the genome-replication inhibitor, GuHCl, (**Figs. 6A, B**). Surprisingly, the signal was the same even when luciferase-coding sequence was present (compare ΔSLII-3 in **Fig. 6B** to ΔIRES in **Fig. 6A**). To determine the limit of detection of the luciferase activity assay, we evaluated the activity of a serial dilution of purified luciferase (**Fig. 6C**). The limit of detection was 1.26 pg of luciferase, corresponding to 10^7^ molecules of luciferase or 1200 molecules of luciferase per cell (**Fig. 6C**). We were unable to achieve a detectable signal by concentrating the sample from ΔSLII-3. Taken together, we conclude that initiation of translation likely occurs downstream of luciferase-coding sequence in non-structural protein- coding sequence.

**Fig 6.**
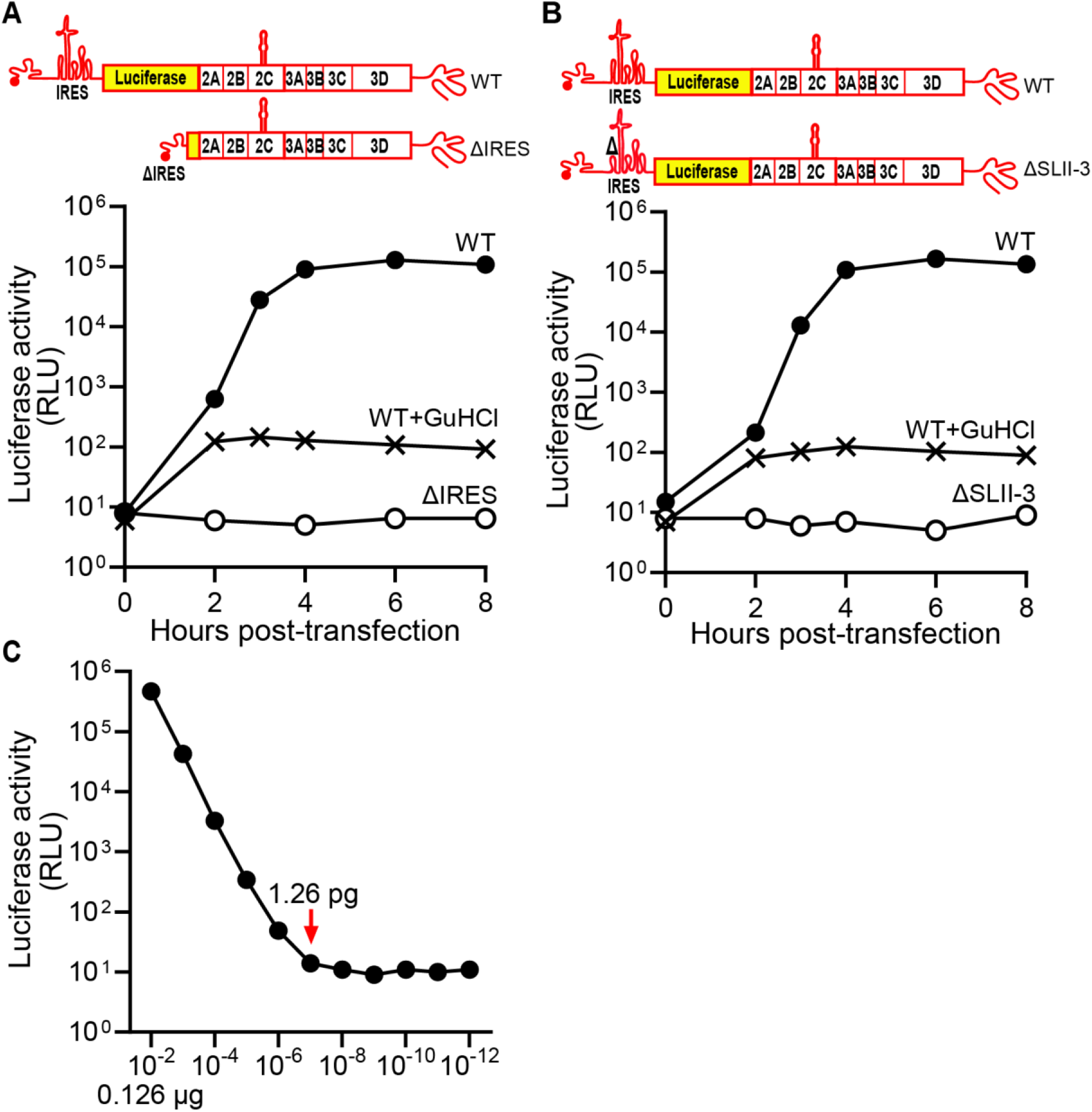
IRES-independent translation may initiate downstream of luciferase coding sequence within the non-structural protein-coding sequence. **(A, B)** Subgenomic replicon luciferase assay comparing the depicted RNAs: WT vs ΔIRES (panel A) and WT vs ΔSLII-3 RNAs (panel B). As a control, the WT RNA was transfected in the presence of guanidine hydrochloride (GuHCl). Luciferase activity is reported in relative light units (RLU) as a function of time post-transfection. Luciferase units for ΔIRES and ΔSLII-3 were not detected above 10^1^. **(C)** Luciferase activity observed for a serial dilution of purified recombinant luciferase enzyme. The initial amount of luciferase in the reaction was 12.6 µg. The limit of detection was reached at a 1 x 10^7^ fold dilution (1.26 pg luciferase), indicated by the red arrow. This corresponds to 1.2 x 10^7^ molecules of luciferase and 1,200 molecules of luciferase per cell from RNAs containing either a deleted IRES or ΔSLII-3.

### Indirect detection of PV 3CD produced by the IRES-independent mechanism

Translation of PV genomes cannot be detected by immunofluorescence in the absence of genome replication [37]. However, the presence of PV 3CD can be detected in the absence of genome replication because of the capacity of this protein in to induce biosynthesis of phosphatidylinositol-4-phosphate (PI4P) [37]. In the uninfected or mock-transfected cell, PI4P localizes to the Golgi apparatus (anti-PI4P in the column marked Mock in **Fig. 7**). Transfection and replication of wild-type PV subgenomic replicon RNA leads to induction and redistribution of PI4P (anti-PI4P in the column marked RLuc-WT in **Fig. 7**). In this circumstance, it is also possible to observe viral proteins (anti-3D in in the column marked RLuc-WT in **Fig. 7**). However, PI4P was also induced in the presence of GuHCl (anti-PI4P in the column marked WT + GuHCl in **Fig. 7**) [37]. Under these conditions, viral proteins were not detected (anti-3D in the column marked WT + GuHCl in **Fig. 7**) [37]. Deletion of the IRES also causes induction of PI4P in the absence (anti-PI4P in the column marked ΔIRES in **Fig. 7**) and presence of GuHCl (anti-PI4P in the column marked ΔIRES + GuHCl in **Fig. 7**). The same is not true for a subgenomic replicon containing stop codons following 3B-coding sequence, which would preclude production of 3CD (anti-PI4P in the column marked 3B STOP in **Fig. 7**). These observations provide support for the expression of 3CD in the absence of the IRES at levels perhaps on par with those expressed by the WT subgenomic replicon in the presence of GuHCl.

**Fig 7.**
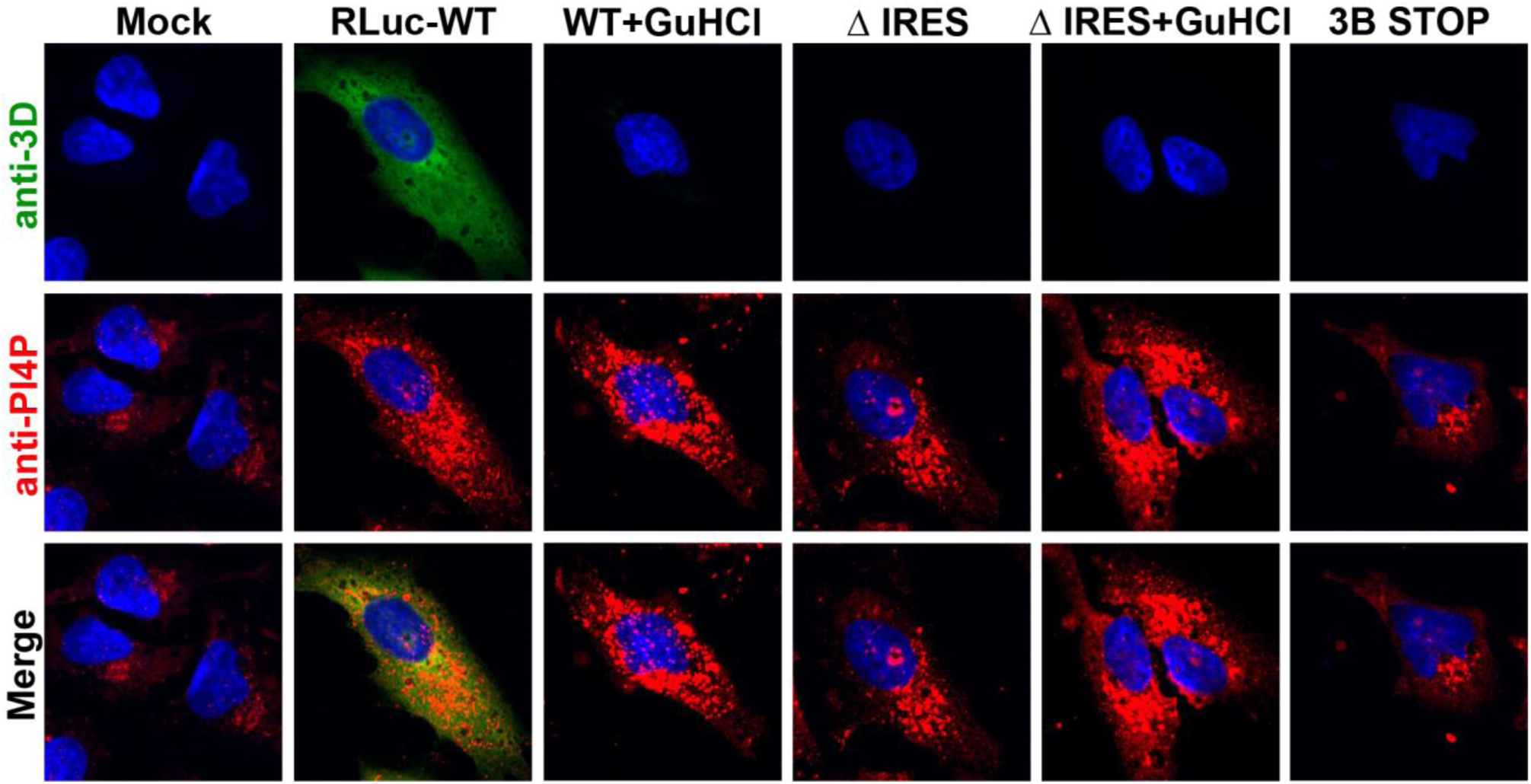
Induction and redistribution of PI4P serves an indirect method to detect production of 3CD by the IRES-independent translation mechanism. Immunofluorescence of cells after transfection. HeLa cells were transfected with *in vitro* transcribed subgenomic replicon RNAs: RLuc-WT; -ΔIRES, and a full-length genomic RNA with two STOP codons after the 3B-coding sequence: 3B STOP. WT and ΔIRES transfected cells were also treated with 3 mM GuHCl. Six hours post-transfection, cells were immunostained for the presence of PI4P (red) and 3D (green). Nucleus was stained with DAPI (blue). Mock represents cells that were taken through the transfection protocol in the absence of RNA. PI4P was induced and redistributed in cells transfected with WT and ΔIRES, both in the absence and presence of GuHCl, but not 3B STOP. 3D was detected in cells transfected with WT in the absence of GuHCl only.

### A primary site of initiation of IRES-independent translation is located within 2A-coding sequence

We evaluated the PV non-structural protein-coding sequence for the presence of methionine residues conserved across members of the enterovirus genus. The first conserved methionine residue was located at the carboxyl terminus of the 2A protease (**Fig. 8A**). We reasoned that if this methionine residue represented the initiating methionine for the IRES-independent mechanism, then recombination should be inhibited by introducing stop codons downstream of the corresponding AUG in the donor genome, which was observed (② х acceptor in **Fig. 8B**). The observed recombination efficiency was reduced by more than 10- fold relative to control (compare to ① х acceptor in **Fig. 8B**). Mutation of this AUG codon to UUG or UUU had no effect on recombination (③ or ④ х acceptor in **Fig. 8B**). However, deletion of this AUG did (⑤ х acceptor in **Fig. 8B**). Previous studies of 2A performed by the Wimmer laboratory demonstrated that the carboxyl terminus of 2A was not important for enzyme function but the nucleotide and/or amino acid sequence in this region was sensitive to mutation [38]. We suggest that the conserved methionine residue is the first residue of the polyprotein produced by the IRES-independent mechanism. However, the sequence of the initiation codon used may not be restricted to AUG codon.

**Fig 8.**
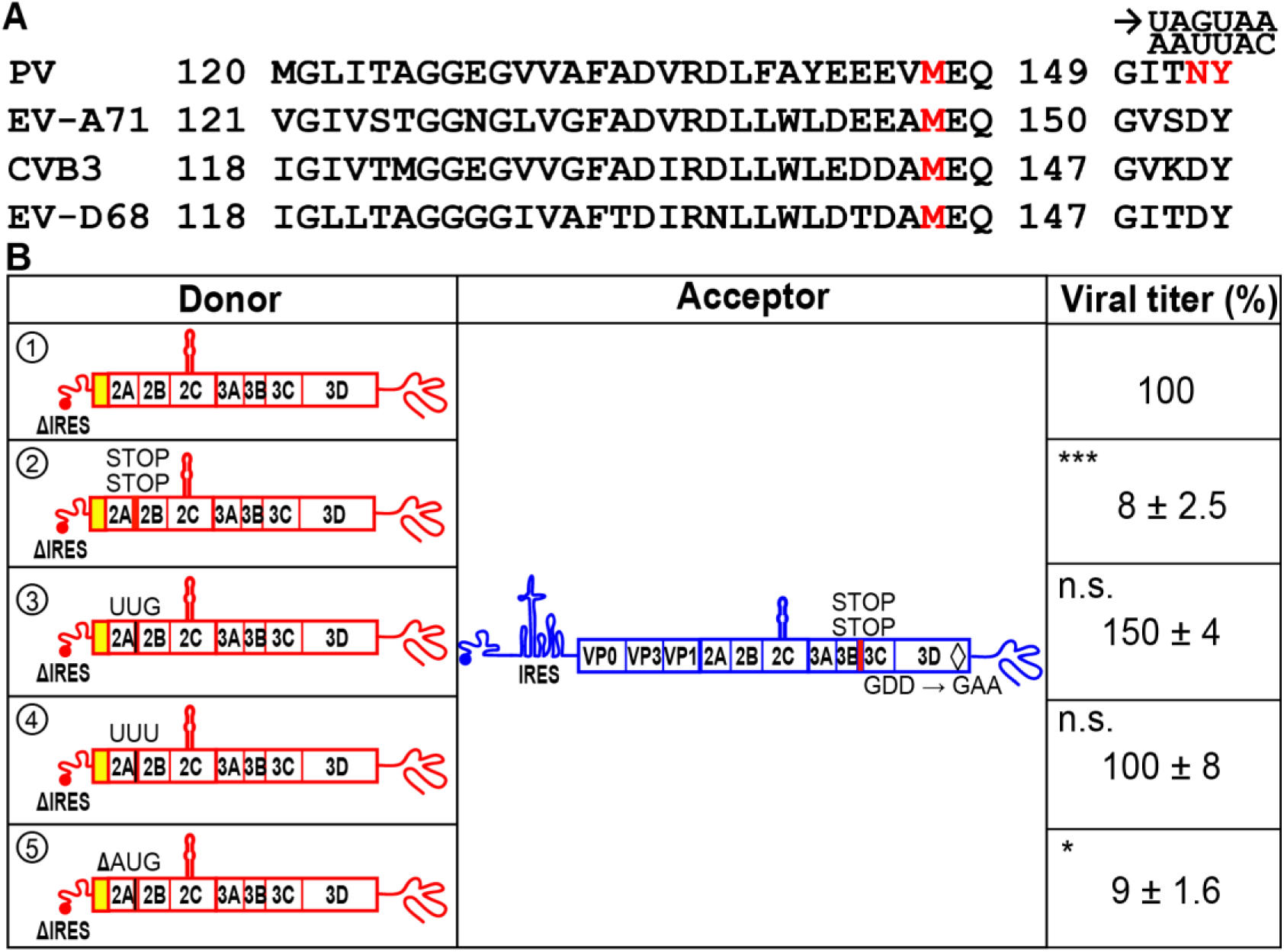
Deletion of a conserved AUG in the 2A-coding sequence reduces translation of the donor RNA leading to a reduction in viral RNA recombinants. **(A)** Primary amino-acid sequence alignment of a portion of 2A sequence from PV, EV-A71, CVB3, and EV-D68. Numbers refer to 2A protein sequence. The conserved methionine is shown in red. The sites for the insertion of two STOP codons are shown in red, the codons AAU and UAC were changed to UAG and UAA, respectively. (**B**) Comparison of infectious virus produced by the indicated donor RNAs with the specified modifications: ①: ΔIRES; ②: Insertion of two STOP codons after the 2A-coding sequence (2A STOP); ③: AUG to UUG; ④: AUG to UUU; ⑤: ΔAUG. Sites for the modifications are depicted. In all cases, the acceptor RNA contained two STOP codons after 3B-coding sequence (3B STOP) and the mutation that inactivates the RdRp (GDD to GAA). Indicated are the relative viral titers with the average viral titer from recombination using ΔIRES donor (①) and acceptor set as 100% (7,500 pfu/mL, mean±SEM; *n = 3*). Statistical analyses were performed using unpaired, two-tailed t-test (*significance level *p* = < 0.05, *** *p* = < 0.001, n.s. = non-significant). The 2A STOP and ΔAUG reduced viral recombinants while the AUG to UUG and AUG to UUU did not.

### Contribution of eIF2A and eIF2D to IRES-independent translation

Initiation of translation in mammals is no longer thought to be as specific as it once was. Use of non-AUG codons under conditions in which eIF2α is active is now well documented [28, 39] (**Fig. 9A**). However, under conditions of stress in which eIF2α is phosphorylated and normal translation is no longer active, alternative mechanisms of translation initiation have evolved [28, 29]. One such mechanism utilizes eIF2A and eIF2D. In addition to initiating at an AUG codon, these alternative initiation factors use a variety of unique and overlapping codons (**Fig.9A**) [28]. The most direct test of the use of eIF2A and eIF2D is genetic ablation of the expression of one or both. This approach was facilitated by using the HAP1 cell line [40, 41]. HAP1 cells are a near-haploid, human cell line derived from the KBM-7 myelogenous leukemia cell line [40]. HAP1 cell lines null for expression of most non-essential genes are available in the Horizon Discovery knockout (KO) collection [42].

**Fig 9.**
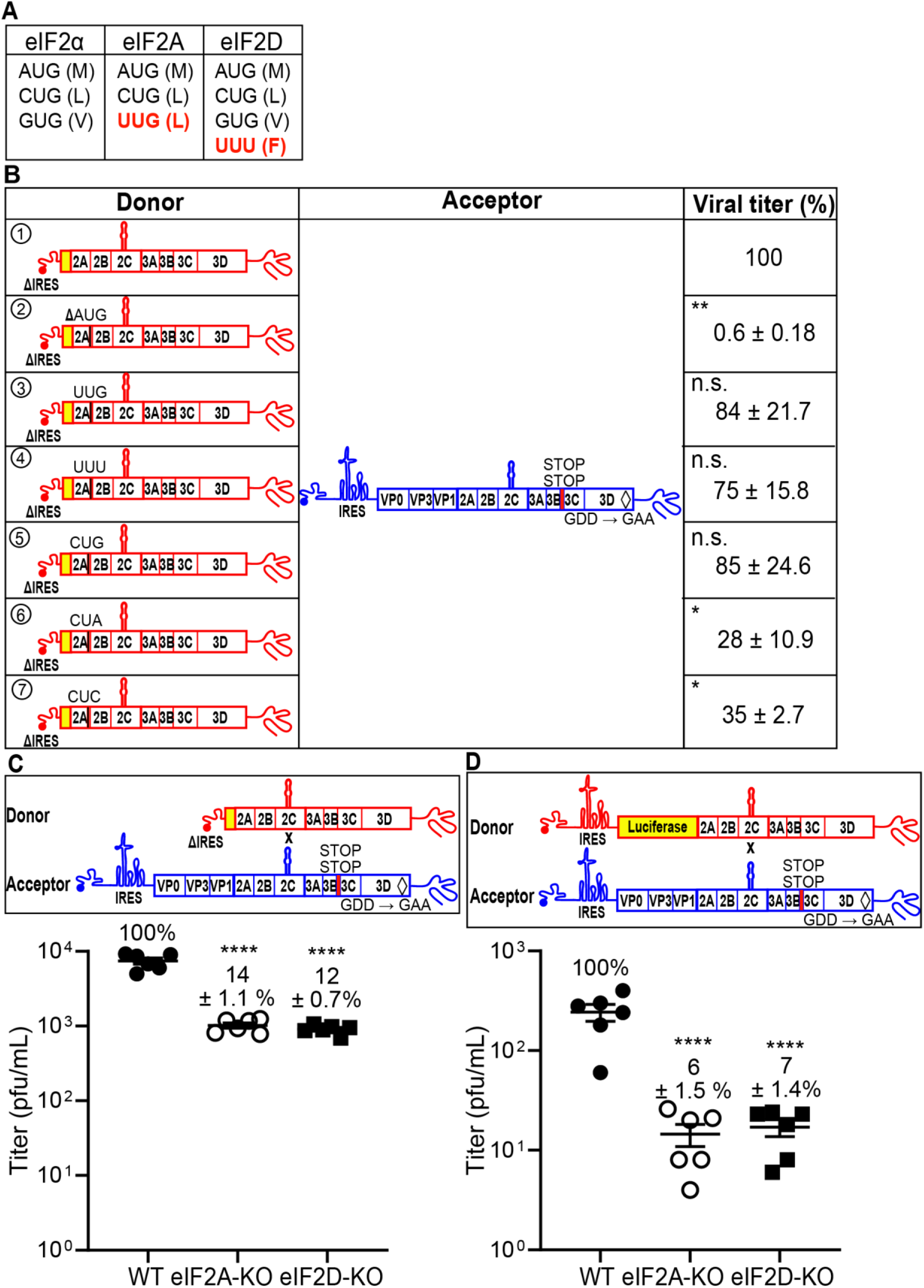
eIF2A and eIF2D initiation factors contribute to IRES-independent translation. **(A)** Start codons utilized by eIF2, eIF2A, and eIF2D initiation factors [28]. Amino acids for each codon is in parentheses. Initiation factor-specific start codons are indicated in red. **(B)** Comparison of infectious virus produced by the indicated donor RNAs with modifications that alter the conserved AUG to initiation factor specific start codons: ①: ΔIRES; ②: ΔAUG; ③: AUG to UUG; ④: AUG to UUU; ⑤: AUG to CUG; ⑥: AUG to CUA; ⑦: AUG to CUC. Codons UUG, UUU and CUG can be utilized by initiation factors but CUA and CUC cannot. In all cases, the acceptor RNA contained two STOP codons after 3B-coding sequence (3B STOP) and the mutation that inactivates the RdRp (GDD to GAA). The indicated set of donor and acceptor RNAs in a 1:5 molar ratio (total 0.3 µg) was co-transfected into HAP1 WT cells. Indicated are the relative viral titers with the average viral titer from recombination using ΔIRES donor (①) and acceptor set as 100% (647 pfu/mL, mean±SEM; *n = 3*). Statistical analyses were performed using unpaired, two-tailed t-test (*significance level *p* = < 0.05, ** *p* = < 0.01, n.s. = non-significant). The ΔAUG, CUA, and CUC reduced viral recombinants significantly while UUG, UUU and CUG did not, consistent with eIF2A and eIF2D contributing to IRES-independent translation. **(C, D)** Viral recombinants are reduced in HAP1 cells deficient for eIF2A and eIF2D expression. The indicated set of donor and acceptor RNAs was co-transfected into HAP1 WT or eIF2A-KO or eIF2D-KO cells. Donor RNA: ΔIRES (panel **C**); RLuc-WT (panel **D**). Relative viral titers with the average viral titer from HAP1 WT set as 100% were shown (panel **C**, 7,433 pfu/mL; panel **D**, 243 pfu/mL; mean±SEM). Statistical analyses were performed using unpaired, two-tailed t-test (****significance level *p* = < 0.0001).

HAP1 cells supported the IRES-independent mechanism of translation in the context of the cell-based PV recombination assay (donor ① in **Fig. 9B**). As we will describe below, these cells also turn out to be susceptible to PV infection. The conserved AUG codon in 2A- coding sequence was required in the HAP1 background (donor ② in **Fig. 9B**). Both unique and shared non-AUG codons used for initiation by eIF2A and eIF2D were well tolerated as substitutions for AUG in the donor genome (donors ③, ④, and ⑤ in **Fig. 9B**). The eIF2A and/or eIF2D-utilized codons, UUG and CUG, both code for leucine. CUA and CUC codons also code for leucine but are not utilized by either eIF2A or eIF2D (**Fig. 9A**) [28]. These two codons were not as well tolerated as the other leucine codons (donors ⑥ and ⑦ in **Fig. 9B**). Together, these data are consistent with a role for eIF2A and/or eIF2D in IRES-independent translation.

To probe the role of eIF2A and eIF2D in IRES-independent translation, we evaluated recombination using the ΔIRES donor in eIF2A-KO and eIF2D-KO cell lines (**Fig. 9C**). The absence of either factor diminished recombination by nearly 8-fold relative to WT cells (**Fig. 9C**). Interestingly, recombination driven by proteins produced by an IRES-dependent mechanism also exhibited a dependence on eIF2A and eIF2D, with recombination reduced by 15-fold in the absence of either factor (**Fig. 9D**).

We hypothesized that the dependence of donor genomes containing a functional IRES on eIF2A or eIF2D might be the result of our transfection of RNA transcribed in vitro. RNA produced in vitro will have a triphosphorylated 5’-end (5’-ppp), a known pathogen associated molecular pattern recognized by several mechanisms of innate immunity [43], including the dsRNA-activated protein kinase, PKR [44]. Transient activation of PKR would lead to phosphorylation of eIF2α and a need for eIF2A or eIF2D. If this is the case, then infection should not exhibit the same dependence on these alternative translation initiation factors.

As a part of our effort to detect IRES-independent translation by monitoring the activity of an expressed reporter, we constructed a genomic replicon harboring nanoLuciferase (NL, Nluc, or nanoLuc) between 2C- and 3A-coding sequence. Surprisingly, this construct actually replicated, yielding a 4-log amplification of nanoLuc activity, and GuHCl inhibited replication (**Fig. 10A**). This tool permitted us to compare outcomes for transfection or infection in the absence or presence of eIF2A or eIF2D. As observed for the traditional luciferase reporter (**Fig. 9D**), replication of this nanoLuc replicon benefited substantially from the presence of eIF2A and eIF2D when RNA was introduced by transfection (**Fig. 10B**). In contrast, introduction of the nanoLuc reporter by infection did not exhibit a significant dependence of either eIF2A or eIF2D (**Fig. 10C**). Simultaneous elimination of both eIF2A and eIF2D exhibited a phenotype equivalent to that observed for the individual knockouts (**Fig. 10D**). We validated the HAP1 KO cell lines used by evaluating expression of eIF2A and eIF2D using Western blotting (**Fig. 10E, F**). Importantly loss of eIF2A or eIF2D did not impact the level of eIF2α in cells (**Fig. 10G**). We were unable to perform the comparable experiments with EV- A71 because this virus does not replicate well or spread in the HAP1 background. We conclude that an IRES-independent, eIF2A/eIF2D-dependent mechanism exists in PV and other enteroviruses to sustain replication under conditions in which the “normal” mechanism for translation initiation is unavailable.

**Fig 10.**
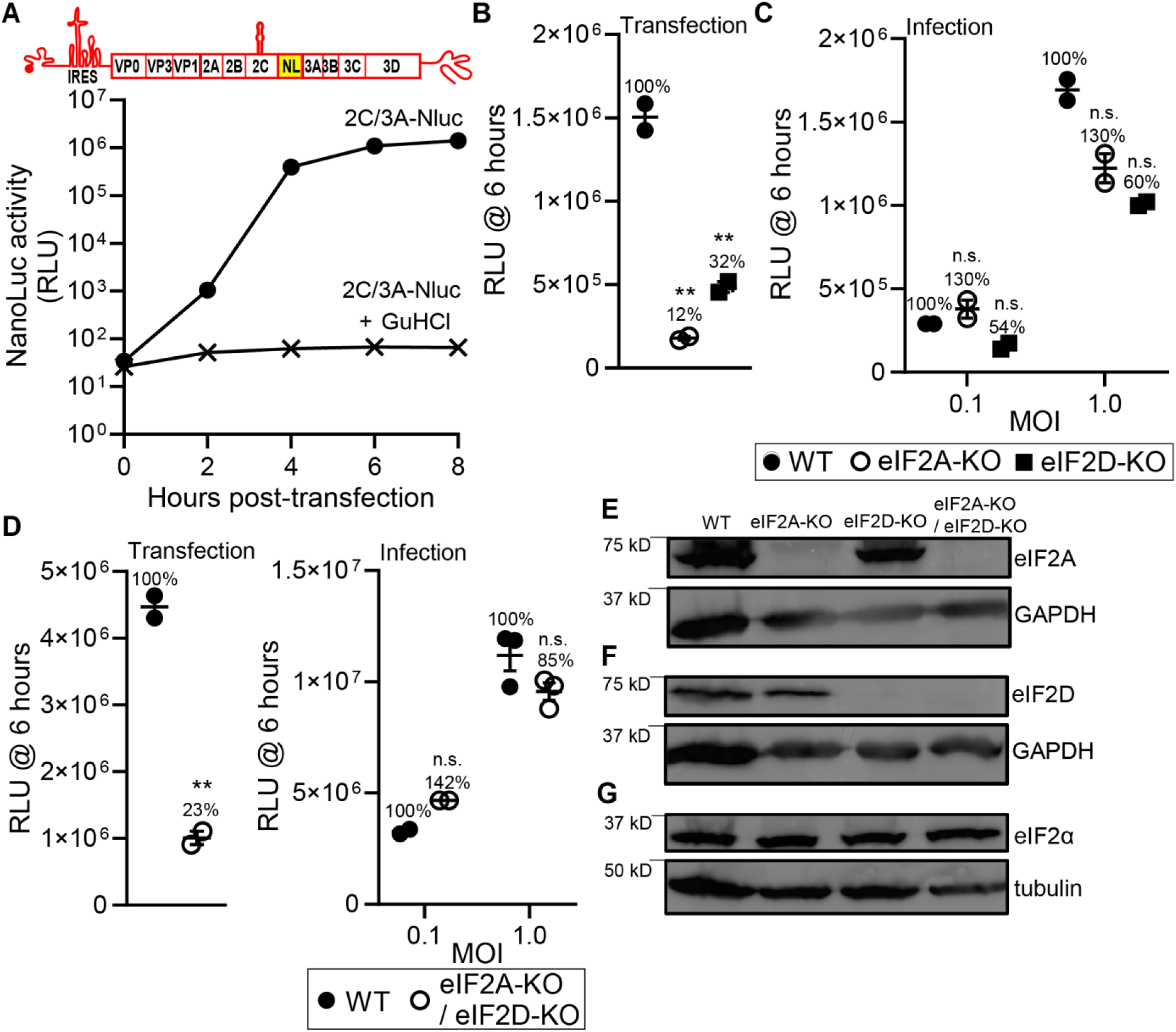
eIF2A and eIF2D contribute to translation of enteroviral RNAs. **(A)** NanoLuc activity in HAP1 WT cells transfected with a full-length PV genome with the nanoLuc-coding sequence embedded between 2C- and 3A-coding region (2C/3A-Nluc). As a control, the PV 2C/3A-Nluc RNA was transfected in the presence of GuHCl. **(B)** Comparison of nanoLuc activity at six hours post-transfection using PV 2C/3A-Nluc RNA in HAP1 WT, eIF2A-KO, and eIF2D-KO cells. Data from one of two biological replicates with similar results, each with two technical replicates. **(C)** Comparison of nanoLuc activity at six hours post-infection using an MOI of 0.1 or 1 in HAP1 WT, eIF2A-KO, and eIF2D-KO cells. Data from one of two biological replicates with similar results, each with two technical replicates. **(D)** Comparison of nanoLuc activity at six hours post-transfection using PV 2C/3A-Nluc RNA (Transfection), and at six hours post-infection using an MOI of 0.1 or 1 (Infection) in HAP1 WT, and eIF2A-KO / eIF2D- KO cells. Data from one of two biological replicates with similar results, each with two or three technical replicates. Statistical analyses were performed using unpaired, two-tailed t-test (**significance level *p* = < 0.01, n.s. = non-significant). **(E-G)** Western blot analysis of eIF2A (panel **E**), eIF2D (panel **F**) and eIF2α (panel **G**) in HAP1 WT, eIF2A-KO, eIF2D-KO, and eIF2A-KO / eIF2D-KO cells. Cells were processed for western blot analysis and probed using anti-eIF2A, eIF2D and eIF2α antibodies. GAPDH and tubulin were used as a loading control for western blot.

### Single-cell analysis suggests a role for eIF2A and eIF2D during infection

It is becoming increasingly clear that analysis of viral replication dynamics at the population level can mask phenotypes [45]. For these experiments, we use a GFP reporter expressed by infectious PV to monitor replication dynamics in 100 – 200 single cells at intervals of 30 min for 12 – 24 h [45, 46]. We evaluate four parameters: (1) start point, which is the time in which the fluorescence becomes detectable and indicates an infection has been established; (2) maximum, which is the maximum value of the observed fluorescence and correlates with the magnitude of genome replication; (3) slope, which is the rate of the fluorescence change and correlates with the speed of genome amplification; and (4) infection time, which is the time from start to maximum and correlates with the virus generation time [45]. We plot distributions of these parameters and perform statistical tests to determine if the perturbation under investigation causes a significant difference relative to the control [45, 46].

Here, we applied this experimental paradigm to determine if a phenotype exists for replication of PV when eIF2A or eIF2D is not present. In the presence of eIF2A and eIF2D, PV establishes infection faster (**Fig. 11A**), replicates to higher levels (**Fig. 11B**), and replicates faster (**Fig. 11C**) than in the absence of one of these factors. The loss of eIF2A or eIF2D essentially prevents replication of PV requiring a longer infection time (**Fig. 11D**). Slow replication may occur in cells better able to use intrinsic defense to delay viral infection by activating eIF2α. The existence of two classes of cells is also supported by evaluation of the percentage of cells infected in the presence or absence of eIF2A/eIF2D. One-quarter of cells exposed to PV do not establish infection in the absence of eIF2A or eIF2D (**Fig. 11E**). Quantitative analysis of the data is presented in **Figs. 11F** and **11G**. We conclude that the IRES-independent, eIF2A/eIF2D-dependent mechanism likely functions during normal infection when certain factors are present or not in the host cell.

**Fig 11.**
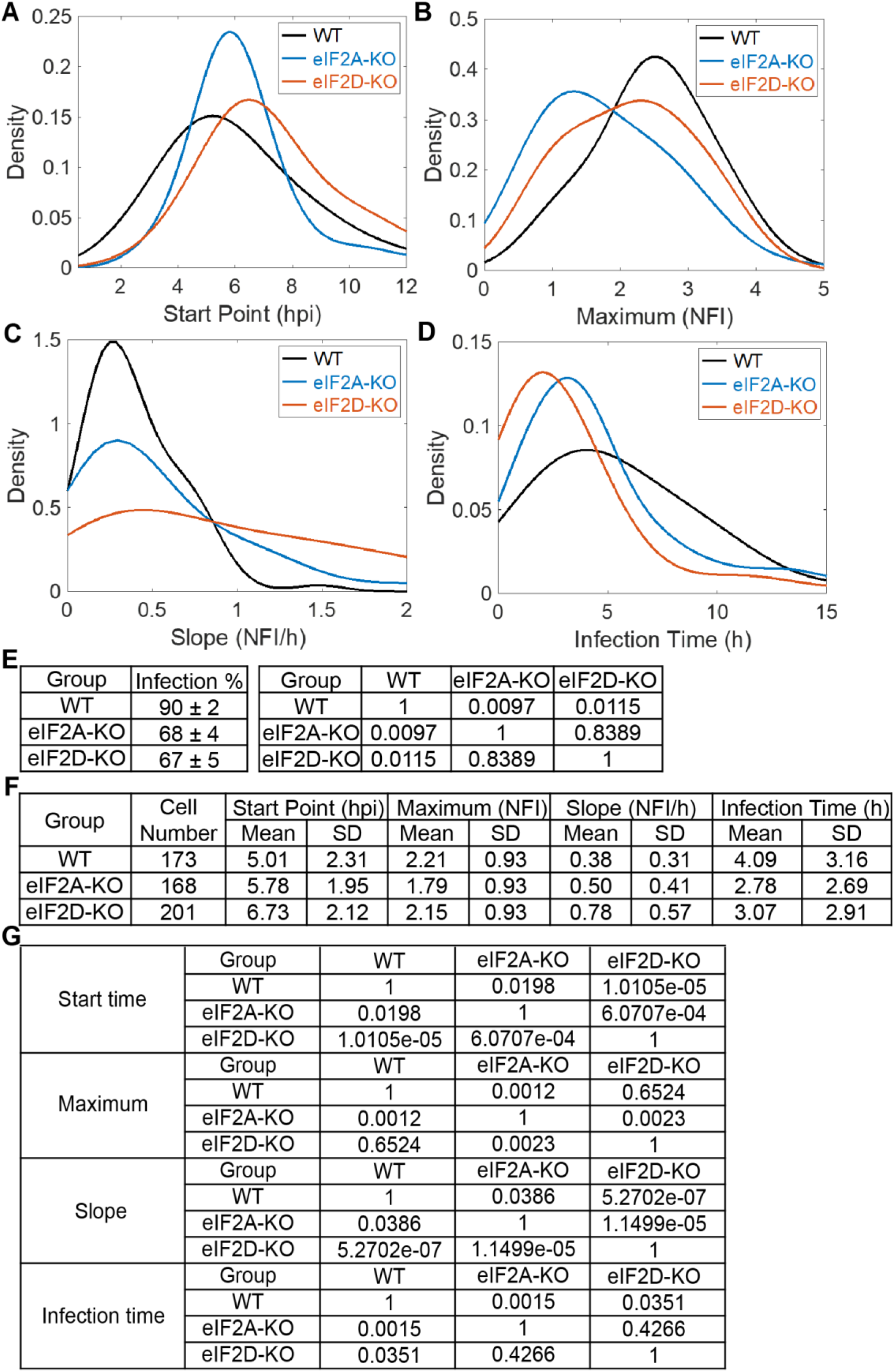
Single-cell analysis suggests a role for eIF2A and eIF2D during infection. Single cell analysis [45, 46] using an MOI of 5 of PV-eGFP_PV_ in HAP1 WT, eIF2A-KO, and eIF2D-KO cells. Comparison of the distributions of each parameter: start point (panel **A**); maximum (panel **B**); slope (panel **C**); and infection time (panel **D**); is shown. hpi, hours post infection; NFI, normalized fluorescence intensity. **(E)** Percentage of PV-infected cells using HAP1 WT, eIF2A-KO, and eIF2D-KO cells (mean±SD, *n*=3) (Left). Adjusted P-values of the t-tests (Right). **(F, G)** Quantitative analysis from the data presented in panels **A**-**D**. Shown are the mean and standard deviation for each of the indicated parameters (panel **F**). Adjusted P- values of the t-tests (panel **G**).

## Discussion

For recombination to occur between two genetically distinct genomes by a template-switching mechanism, the two genomes must be localized to the same genome-replication organelle. How enteroviral genomes are trafficked to the site of replication, however, is not known. The central hypothesis driving this study was that determinants located in the polyprotein mediated localization to the genome-replication organelle, perhaps co-translationally (**Figs. 1A, B**). Therefore, translation of both genomes would be a prerequisite for recombination (**Figs. 1C, D**). Such a requirement would be consistent with previous studies by other laboratories that have established a requirement for translation of the polyprotein in cis for genome replication to occur [34, 35]. However, we observed that inactivation of translation of a donor PV genome by deleting the IRES enhanced recombination by 5-fold over control (**Fig. 2**).

Studies of several positive-strand RNA viruses, including PV, have suggested the existence of non-replicative recombination [27, 47, 48]. Such a mechanism is independent of the RdRp and occurs by a direct chemical ligation of two overlapping fragments of RNA [49]. However, our studies demonstrated the requirement for an active RdRp in the IRES-deleted, donor PV genome (**Fig. 3**). The dependence on an active RdRp would, by definition, indicate replicative recombination and necessitate translation by an IRES-independent mechanism. Because the IRES eliminated the entirety of the 5’-UTR, it was possible that the normal constraints on translation initiation were lost, and we were observing something cryptic and/or irrelevant [50]. We ruled out this possibility by inactivating the IRES with a 10-nt deletion; this donor PV genome also supported recombination in an RdRp-dependent manner (**Fig. 4**). We reached similar conclusions using EV-A71 (**Fig. 5**). Looking back at all of the studies supporting non-replicative recombination, they all employed RNA transfection of RNAs capable of expressing the RdRp using an IRES-independent mechanism [13, 27]. These previous studies never considered an alternative translation mechanism.

Together, our results suggested the compelling possibility that an IRES-independent mechanism exists for translation of the non-structural region of the polyprotein. We were unable to detect viral proteins directly or indirectly by using standard reporters (**Fig. 6**). However, we were able to exploit our previous observation that 3CD induces PI4P, even when present at levels undetectable by immunofluorescence [37]. Constructs lacking the IRES induced PI4P in a manner requiring translation through the 3C- and 3D-coding sequence (**Fig. 7**). We identified an AUG codon in the 3’-end of 2A-coding sequence that is conserved among viruses comprising the enterovirus genus (**Fig. 8A**). The Wimmer lab showed years ago that this region of the protein was not essential for protease function but suggested the existence of cis-acting replication element in this region [38]. Our reverse- genetic analysis supported the use of this AUG as the start site for IRES-independent translation (**Figs. 8** and **9**).

What was unexpected, however, was that multiple, non-AUG codons also supported translation initiation (**Figs. 8** and **9**). Even more unexpected was that translation initiation depended on eIF2A and eIF2D (**Fig. 9**). These factors are essential for translation under conditions in which eIF2α has been phosphorylated, for example in response to activation of a mediator of the integrated stress response [29]. These factors also use non-AUG codons for initiation (**Fig. 9A**) [28]. However, the dependence on eIF2A and eIF2D was observed not only for a genome lacking an IRES (**Fig. 9C**) but also for a genome containing an IRES (**Fig. 9D**), yet another unexpected result. All of our experiments launched infection by transfection of RNA produced in vitro using T7 RNA polymerase. We did not observe a strong dependence on eIF2A and/or eIF2D when infection was launched using virus (**Fig. 10**).

One explanation for a requirement for eIF2A and/or eIF2D when in vitro transcribed genomes are used is that these genomes likely activate intrinsic antiviral defenses that normal infection would not. RNA produced by in vitro transcription contains a triphosphate at the 5’-end while the authentic viral genome would contain the 3B-encoded, genome-linked protein (VPg) [51]. The 5’-triphosphate is a well-known pathogen associate molecular pattern recognized by antiviral pattern recognition receptors like the double-stranded RNA-activated protein kinase (PKR) [44], the interferon-inducible protein with tetratricopeptide repeats (IFIT) [52], and the retinoic acid-inducible gene product (RIG-I) [53]. Therefore, the IRES-independent, eIF2A/eIF2D-dependent mechanism of translation may have evolved to antagonize viral restriction by eIF2α kinases and IFITs.

Initiation of translation on the PV IRES uses several cellular factors [54]. eIF4G and eIF4A bind to the IRES and recruit the 43S preinitiation complex comprised of the 40S ribosomal subunit, eIF3, tRNA-Met_i_-bound eIF2, among other factors (**Fig. 12A**) [55]. We are not the first to suggest the use of an alternative translation mechanism during PV infection. The Lloyd laboratory showed that translation of the PV polyprotein late in infection does not require eIF2 [56]. The best-known eIF2-less mechanisms replace eIF2 with eIF5B [57, 58] or MCT- 1•DENR (multiple copies in T-cell lymphoma • density regulated protein complex) (**Fig. 12B**) [59, 60]. Replacement of eIF2 eliminates sensitivity to eIF2α kinases and expands the possibility for use of non-AUG codons. However, the components and mechanism for assembly of the preinitiation complex are thought to be unchanged relative to the eIF2- dependent mechanism (**Figs. 12A,B**) [54].

**Fig 12.**
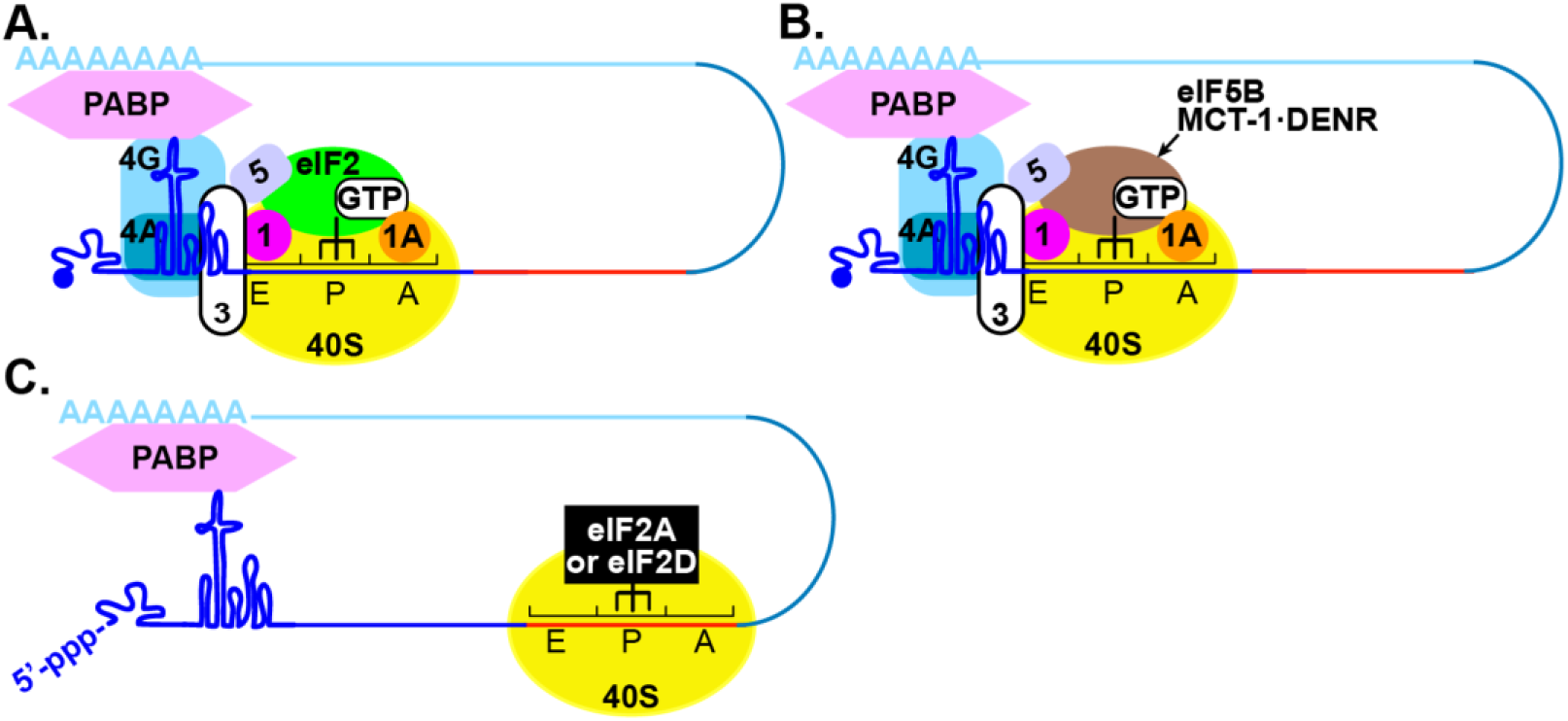
Models for initiation of translation on the enterovirus genome. **(A)** Under normal conditions, eIF4G and eIF4A bind to the IRES and recruit the 43S preinitiation complex composed of the 40S ribosomal subunit (yellow), eIF2/GTP/Met-tRNAi^Met^ (green) and eIF3, among other factors. Initiation may also be facilitated by interactions of the poly(rA)-binding protein, PABP, bound to the 3’-poly(rA) tail. This interaction may use eIF4G or viral/cellular factors interacting with the cloverleaf. Translation begins at the AUG start site in an eIF2- directed manner. **(B)** Multiple distinct eIF2-independent mechanisms exist. Both eIF5B or MCT-1•DENR (brown) can substitute for eIF2 to promote recruitment of initiator tRNA and translation initiation at the AUG start site. These factors may also initiate at non-AUG codons. **(C)** An IRES-independent, eIF2A/eIF2D-dependent mechanism. Activation of intrinsic antiviral defense mechanisms, for example as a result of the presence of 5’-ppp on our biosynthetic enteroviral genomes, will lead to inactivation of eIF2 and sequestration of eIF3. The non- canonical translation initiation factors eIF2A and eIF2D (black) direct translation initiation from a region of RNA (red) within 2A-coding sequence in a manner that is not dependent on the presence of an AUG codon. Other than the 40S ribosomal subunit, factors leading to formation of the translation-initiation complex are not known.

Essentially nothing is known about the eIF2A/eIF2D-dependent mechanism(s) of translation initiation in cells (**Fig. 12C**) [30]. However, this mechanism is essential for cells to survive under stress and has been linked to neurodegenerative diseases and cancer [29]. Efforts to demonstrate a role for eIF2A and/or eIF2D in viral translation have been inconclusive [32, 33, 61, 62], with the majority view being the absence of a role for either factor [61, 62]. Therefore, our demonstration that enteroviruses may use these factors when intrinsic antiviral defenses have been activated may represent an opportunity not only to pursue a role for these factors in other viral systems but also to begin to define the cis-acting RNA determinants driving initiation of translation using these factors. As discussed above, numerous mechanisms exist to function in the absence of active eIF2, so the eIF2A/eIF2D- dependent mechanism may further extend independence from canonical initiation factors. One candidate would be eIF3. IFITs activated by 5’-ppp-RNA sequester eIF3 to diminish translation. The 5’-ppp-PV genomes used here should activate IFITs. The IRES-independent, eIF2A/eIF2D-dependent mechanism would therefore be predicted to function in the absence of eIF3.

Our studies of PV infection at the population level concealed any significant difference in replication in the presence or absence of eIF2A or eIF2D (**Figs. 10C, D**). However, studies at the single-cell level revealed a clear advantage to having both genes expressed (**Figs. 11A, B**). There were two intriguing observations. First, infections that required longer than 5 h for completion appeared to be lost in the absence of eIF2A or eIF2D (**Fig. 11E**). This population may reflect those in which the IRES-independent, eIF2A/eIF2D-dependent mechanism was required, perhaps because the infecting PV variant activated intrinsic defenses. Second, 25% of the cells appeared recalcitrant to infection in the absence of eIF2A or eIF2D (**Fig. 11E**). Perhaps intrinsic defenses are upregulated independent of infection in this population of cells. The bottom line is that virus variant- and/or host-driven circumstances exist that require eIF2A and/or eIF2D for proper resolution, and the requirement for these factors may only become apparent by using a single-cell analysis.

The literature on eIF2A and eIF2D suggests that each functions independently of the other. This circumstance may merely reflect that each protein has been studied in the absence of the other. Our results show quite clearly that loss of eIF2A or eIF2D alone reduces translation to the same extent and causes an equivalent phenotype (**Fig. 10B**), without any further reduction observed when expression of both proteins is eliminated (**Fig. 10D**). These data suggest cooperation between these proteins. The stability of each is independent of the other (**Fig. 10D**). Perhaps recruitment to or function on the ribosome requires both when eIF2A/eIF2D-dependent mechanism of translation is engaged.

Group B enteroviruses establish persistent infections in the heart and pancreas that lead to myocarditis and diabetes, respectively [63, 64]. Deletion of the 5’-terminal cis-acting replication element, the so-called cloverleaf, is a signature of the genomes that persist in the heart [65]. How these deletions impact IRES-dependent translation is unclear, but viral proteins are below the limit of detection during persistent infection [65]. The IRES-independent, eIF2A/eIF2D-dependent translation mechanism exhibits all of the features one might expect to support persistence of an enteroviral genome. Capsid proteins would not be made; so particles would not be released, leading to inflammation or immune responses. The 2A protease is quite efficient at impairing host cell functions [66], and this protein would not be made. The 3C(D) protease can also be toxic [67]. Much of the damage caused by this protein happens late in infection, so there is likely a requirement for high concentrations of the protein. None of the non-structural proteins accumulates to detectable levels. Genomes dependent on this alternative translation mechanism replicate and recombine as demonstrated herein.

In conclusion, this study has revealed an eIF2A/eIF2D-dependent mechanism of translation initiation used by PV that may be conserved in all enteroviruses that appears to permit continued translation and genome replication under conditions in which intrinsic antiviral defenses have been activated. Discovery of the cis-acting determinants driving this alternative translation mechanism will further illuminate the contributions of this mechanism to viral multiplication, fitness, and pathogenesis. Finally, the existence of an mRNA sequence capable of directing eIF2A/eIF2D-dependent initiation of translation may enable elucidation of the repertoire of factors and pathway used for initiation.

## Materials and Methods

### Cells and Viruses

Adherent monolayers of HeLa and L929 fibroblasts were grown in DMEM/F-12 media. Adherent monolayers of human embryonic rhabdomyosarcoma (RD) were grown in DMEM media. Wild-type HAP1 human haploid cells and HAP1 cells knocked-out (KO) for eIF2A (cat# HZGHC002650c001) or eIF2D (cat# HZGHC002651c012) were purchased from Horizon Discovery Group plc. All media was supplemented with 100 U/mL penicillin, 100 µg/mL streptomycin, and 10% heat inactivated (HI)-FBS. All cells were passaged in the presence of trypsin-EDTA.

Wild-type and recombinant viruses were recovered following transfection of RNA produced *in vitro* (see below) from full-length cDNA or from the cell-based assay parental partners. PV type 1 (Mahoney) and EV-A71 (C2) were used throughout this study. Virus was quantified by pfu/mL. Where stated, guanidine hydrochloride (Sigma) was added to growth media at 3 mM.

### Plasmids, in vitro transcription, cell transfection, and virus quantification

Subgenomic PV replicon, ΔIRES, and a replication-incompetent full-length PV genomic RNA were previously described [13]. All insertions / deletions were created using either overlap extension PCR or gBlock gene fragments from IDT. Oligonucleotides used in this study can be found in **S1 Table**. The presence of the desired insertions / deletions, and the absence of additional mutations were verified by DNA sequencing. 3B STOP was modified from a full-length PV genomic RNA by introducing two STOP codons (UAGUAA) after the 3B-coding sequence. ΔSLII-3 was modified from a subgenomic PV replicon by introducing a ten-nucleotide deletion (nt 185-189, nt 198-202) into the SLII-3 region of the IRES [36]. For a full-length PV genome with the nanoLuc-coding sequence embedded between 2C- and 3A-coding region (2C/3A-Nluc), nanoLuciferase-encoding sequence carrying a 3C protease cleavage site at its carboxyl terminus was inserted between the 2C and the 3A region of the PV sequence. Translation occurred from the natural poliovirus initiation codon. Proteolytic cleavage and release of nanoLuciferase occurred by normal 3C protease activity. Plasmids encoding PV genomes (full length or subgenomic) were linearized with *ApaI*. The EV-A71 C2 replicon and ΔIRES were previously described [15]. 3B STOP was modified from a previously described EV-A71 C2-MP4 infectious clone [15, 68]. The EV-A71 C2 replicon and ΔIRES were linearized with *SalI*. 3B STOP was linearized with *EagI*.

All linearized cDNAs were transcribed *in vitro* using T7 RNA Polymerase treated with 2U DNAse Turbo (ThermoFisher) to remove residual DNA template. The RNA transcripts were purified using RNeasy Mini Kit (Qiagen) before spectrophotometric quantification. Purified RNA in RNase-free H_2_O was transfected into cells using TransMessenger (Qiagen). Virus yield was quantified by plaque assay in either HeLa (PV) or RD (EV-A71) cells. Briefly, cells and media were harvested at time-points post transfection (specified in main text or below), subjected to three freeze-thaw cycles, and clarified. Supernatant was then used on fresh cells in 6-well plates; virus infection was allowed to continue for 30 min. Media was then removed, and cells were washed with PBS (pH 7.4) before a 1% (w/v) agarose-media overlay was added. Cells were incubated for either 2-3 days for PV or 3-4 days for EV-A71 and then fixed and stained with crystal violet for virus quantification.

### Antibodies

Antibody for staining PI4P was purchased from Echelon Biosciences. Antibody against PV 3D was produced in Cameron lab. Mouse monoclonal anti-GAPDH antibody (10R- G109a) was purchased from Fitzgerald Industries. All secondary antibodies used for immunofluorescence were purchased from Invitrogen. Secondary antibodies for western blotting were purchased from Amersham GE Healthcare (rabbit anti-HRP) and Cell Signaling (mouse anti-HRP). Rabbit polyclonal anti-eIF2D antibody (12840-1-AP) and anti-eIF2A antibody (11233-1-AP) were purchased from Proteintech. Rabbit polyclonal anti-eIF2α (9722) was purchased from Cell Signaling Technology Inc.

### Indirect immunofluorescence

HeLa cells were fixed 6 hours post-transfection using 4% formaldehyde in PBS for 20 min, followed by washing in PBS and permeabilizing with 20 μM digitonin for 10 min. Digitonin was removed, and cells were rinsed three times with PBS. Cells were then blocked with 3% BSA in PBS for 1 hour and incubated with primary antibodies for 1 hour. Following washes, cells were incubated with secondary antibodies for 1 hour, followed by a 10 min DAPI incubation. The processed coverslips were mounted on glass slides using ProLong Glass Antifade Mountant (Thermo Scientific). Samples were imaged using the Keyence BZ-X800. Primary antibodies: anti-PI4P (1:200); and anti-3D (1:100). All dilutions were made in the blocking buffer.

### Western blotting

Cells were lysed in radioimmunoprecipitation assay (RIPA) buffer with protease inhibitor and clarified by centrifugation. The lysate was mixed with 4× Laemmli buffer, boiled, and then processed by SDS-PAGE. Proteins were transferred to a polyvinylidene fluoride membrane by semi-dry transfer using the Transblot Turbo apparatus (BioRad). Membranes were blocked in 5% dry milk with 0.1% Tween-20, and probed with anti-eIF2A (1:1000), anti-eIF2D (1:1000), and anti- eIF2α (1:1000) antibodies overnight. Anti-rabbit / mouse immunoglobulin G antibody coupled to peroxidase (Amersham GE Healthcare) were used as secondary antibody at a 1:5000 dilution. Protein bands were visualized with the ECL detection system (Amersham, GE Healthcare). Where stated, membranes were subjected to re-probing against anti-tubulin (1:5000) or anti-GAPDH (1:5000).

### Luciferase assays

Subgenomic luciferase assays were performed as described previously [21]. For each time-point, 1.0 x 10^5^ cells were suspended in 100 µl lysis buffer and 10µl was used for measuring luciferase activity. Considering the corresponding luciferase molecules to the limit of detection, up to 1,200 molecules (1.2 x 10^7^ molecules / 1.0 x 10^4^ cells) of luciferase may exist in a cell transfected with RNAs containing either a deleted IRES or ΔSLII-3.

NanoLuc assays were carried out with the NLuc GLOW Assay kit (Nanolight Technology, #325) following the manufacturer’s protocol.

### Cell-based recombination assay

Assays were used as described previously [13–15, 69]. For the cell-based assay using HAP1 cells, unless otherwise stated, equivalent ratios of RNA (total 0.5 µg) were co-transfected into HAP1 cells in 12 well dishes. Cells and media were harvested at 6 hours post transfection, subjected to three freeze-thaw cycles, and clarified. Virus yield was quantified by plaque assay in HeLa cells.

### Single cell analysis

For each group, cells were infected with PV-eGFP_PV_ at the MOI of 5 PFU per cell. On-chip experiments and data processing were done as described previously [45, 46].

---

**S1 Table.**
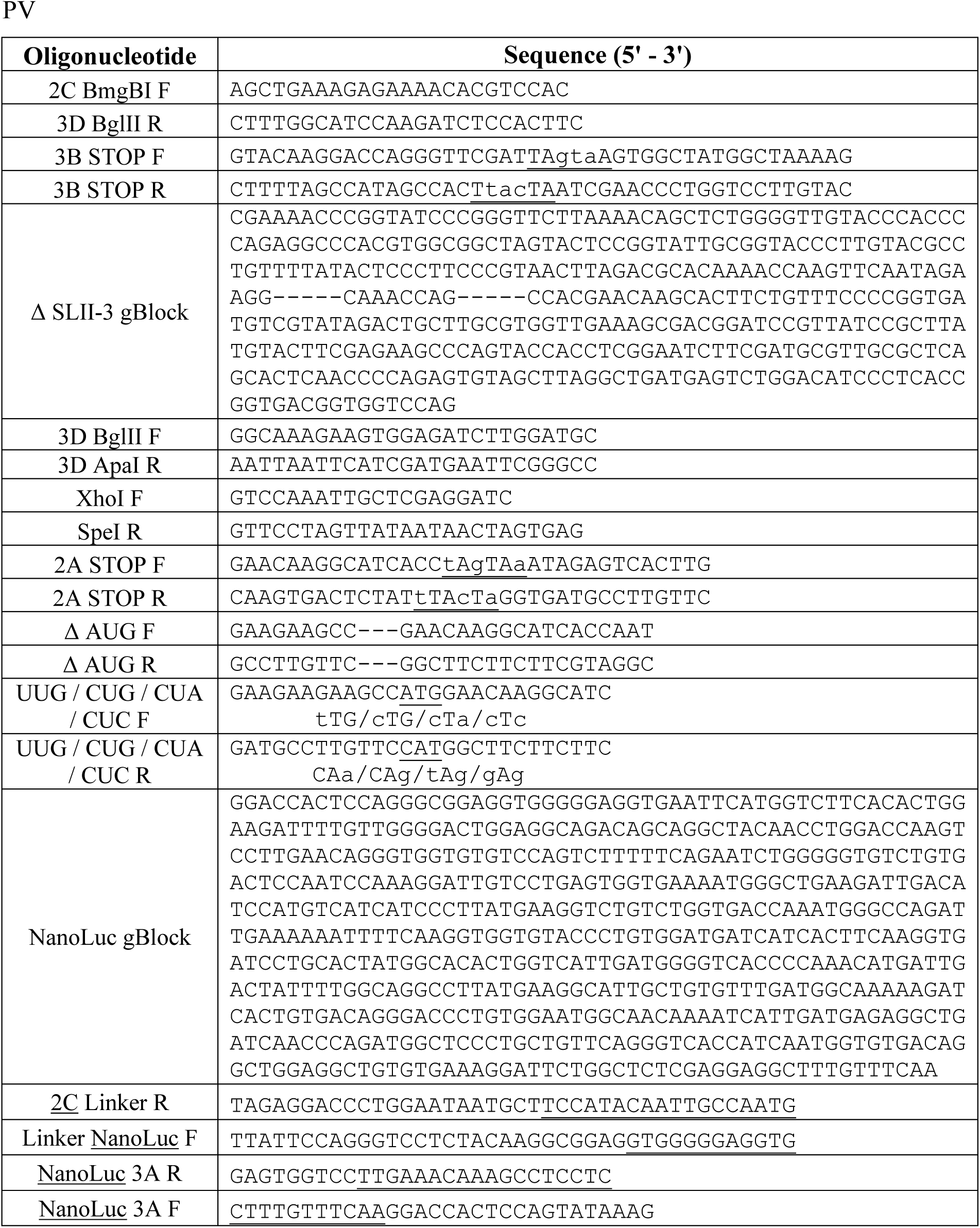
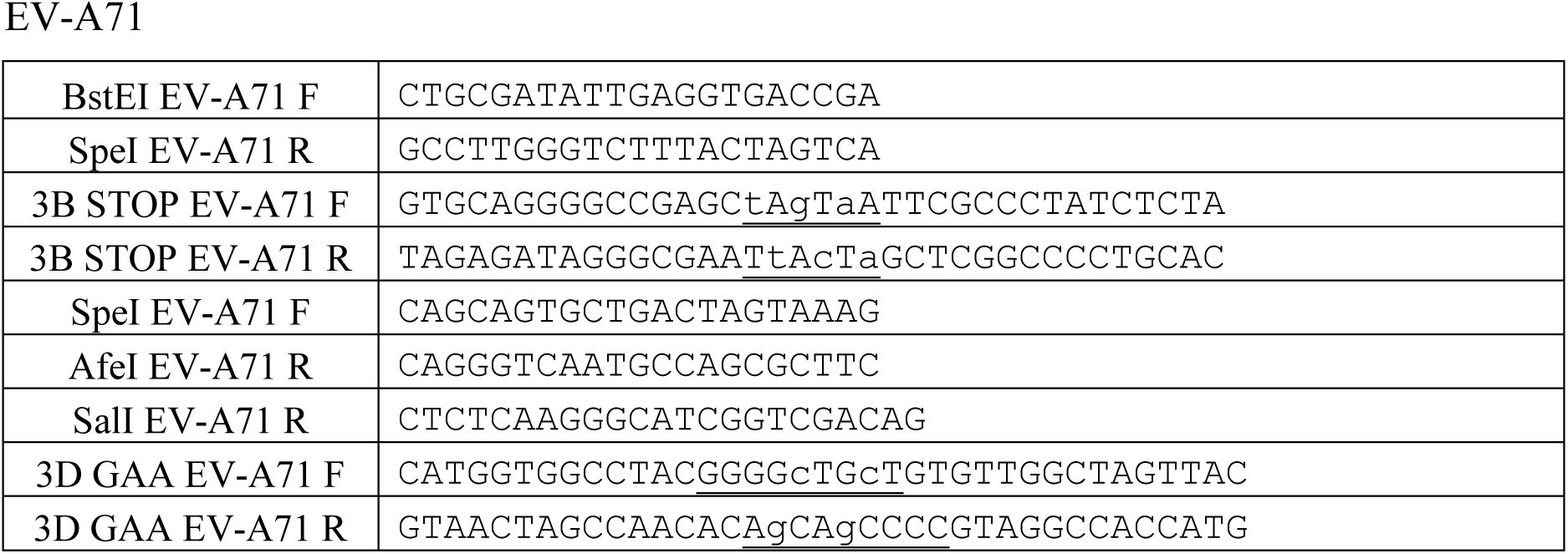
Oligonucleotides used in the study

